# Optical Neuroimage Studio (OptiNiSt): intuitive, scalable, extendable framework for optical neuroimage data analysis

**DOI:** 10.1101/2024.09.17.613603

**Authors:** Yukako Yamane, Yuzhe Li, Keita Matsumoto, Ryota Kanai, Miles Desforges, Carlos Enrique Gutierrez, Kenji Doya

## Abstract

Advancements in calcium indicators and optical techniques have made optical neural recording a common tool in neuroscience. As the volume of optical neural recording data grows, streamlining the data analysis pipelines for image preprocessing, signal extraction, and subsequent neural activity analyses becomes essential. There are a number of challenges in optical neural data analysis. 1) The quality of original and processed data needs to be carefully examined at each step. 2) As there are numerous image preprocessing, cell extraction, and activity analysis algorithms, each with pros and cons, experimenters need to implement or install them to compare and select optimal methods and parameters for each step of processing. 3) To ensure the reproducibility of the research, each analysis step needs to be recorded in a systematic way. 4) For data sharing and meta-analyses, adoption of standard data formats and processing protocols is required. To address these challenges, we developed Optical Neuroimage Studio (OptiNiSt) (https://github.com/oist/optinist), a framework for intuitively creating calcium data analysis pipelines that are scalable, extendable, and reproducible. OptiNiSt includes the following features. 1) Researchers can easily create analysis pipelines by selecting multiple processing modules, tuning their parameters, and visualizing the results at each step through a graphic user interface in a web browser. 2) In addition to common analytical tools that are pre-installed, new analysis algorithms in Python can be easily added. 3) Once a processing pipeline is designed, the entire workflow with its modules and parameters are stored in a YAML file, which makes the pipeline reproducible and deployable on high-performance computing clusters. 4) OptiNiSt can read image data in a variety of file formats and store the analysis results in NWB (Neurodata Without Borders), a standard data format for data sharing. We expect that this framework will be helpful in standardizing optical neural data analysis protocols.

## Introduction

Given the increasing adoption of two-photon, light-sheet, and endoscopic microscopes, huge volumes of optical imaging data are being collected. Simultaneously, various image processing and cell detection software tools have been developed and are available in open source [1–4]. Additionally, a wide variety of downstream analyses for massive cell populations have been published, and many are available in open source. Using the same dataset can yield different analysis results depending on the algorithm employed. For instance, in tasks, such as cell detection or spike inference from fluorescence dynamics, validating the algorithm by obtaining ground truth data is often impractical [5]. Misclassifying or neglecting the neural signals might affect experimental results [6–8]. Determining which algorithm is preferable for a given dataset is challenging. Ideally, it is necessary to employ multiple algorithms for comparison and evaluation [9]. However, as individual algorithms are provided for specific environments (e.g., versions of operating system, programming language, and libraries), researchers who wish to apply new methods to their dataset and compare them with other methods need to prepare different environments for different analytical tools and sometimes reshape their data to fit each tool. They also need to visualize the analysis results, often by writing codes themselves. Additionally, the application of some algorithms needs extensive computation resources and high-performance computer (HPC) clusters to obtain results within a reasonable amount of time [10].

Therefore, we developed a comprehensive framework, the Optical Neuroimage Studio (OptiNiSt), which helps researchers try multiple data analytical methods quickly, interactively visualize the results, and construct data analysis pipelines for reproducible analyses and efficient processing on HPC clusters. OptiNiSt allows users to test multiple analytical methods with different parameters without having to write codes or set up computing environments. The workflow pipeline can be saved coherently with the analysis results for better reproducibility. OptiNiSt aims to make a wide variety of analytical methods accessible to experimental researchers, to promote the use of new analytical methods proposed by theoretical researchers, and at the same time, to promote the standardization of analysis protocols. OptiNiSt’s saving format is compliant with Neurodata Without Borders (NWB) standards [11], resulting in widespread usage because of the diligence in maintaining the community. OptiNiSt is free and open source, uses the GNU General Public License v3.0, and is hosted on GitHub.

There has been a consensus that the creation of an open-source neuroscience database is essential. To achieve Findable, Accessible, Interoperable, and Reusable (FAIR) data [12], sharing not only original data but also relevant information, such as experimental protocols, source code, and research software used for processing or analyzing these datasets, is essential for reproducing research findings [13]. While open science principles are largely accepted, it is practically difficult for researchers to prioritize effective research data management over other processing demands [9]. Due to the complexity and diversity of data structures and lack of established protocols or formats in analytical software, complete universality is still not anticipated, and there are currently no available data or databases that many researchers can reuse or reanalyze easily. To efficiently share data and analysis pipelines among researchers, a framework has to automate the process of saving analysis workflows and results in a standardized format. Reproducibility can be increased by using a comprehensive data analysis pipeline framework. Additionally, researchers who develop data analytical methods need their proposed tools to be tested and used by many researchers to contribute to the community and improve their methods. By including their analytical methods in a common framework, they are more easily accessible and likely to be used. Thus, a comprehensive framework for data analysis pipelines is also beneficial for those who develop data analytical methods.

There are some software applications for calcium imaging data analyses that can compare different analytical methods. CIAtah [14] is a specialized tool for preprocessing and cell detection in calcium imaging data analysis and has the ability to compare different algorithms, but it is implemented in MATLAB, and therefore, only licensed MATLAB users can use it; moreover, it does not save a complete record of analysis pipelines. Mesmerize [15] is written fully in Python, making it readily available, but the desktop graphical user interface (GUI) is not supported any more. Thus, these frameworks are still not comprehensive enough for use as a standard framework for FAIR data sharing to meet the broad demands of calcium imaging data analyses.

## Design and implementation

OptiNiSt was developed with four major design principles: 1) intuitive GUI, 2) extendable to include a variety of analysis modules, 3) reproducible and scalable to be deployed on HPC clusters, and 4) adoption of standard formats for interoperability. The users interact with OptiNiSt through a web browser as the frontend. OptiNiSt has three interfaces, WORKFLOW, VISUALIZE, and RECORD. WORKFLOW is for creating analysis pipelines, VISUALIZE is for visualizing plots of the analysis results, and RECORD is for managing the created pipelines. Through these three GUIs, the user can create and run the analysis pipelines and inspect the results without coding any scripts. All the information necessary to reproduce the pipeline, including the path to the original data and parameters for analyses, is saved in a folder with a specific ID, and the list of pipelines is easily recognized and reproduced through the RECORD interface. The system is also designed to transfer pipelines to HPC clusters for script execution. The analysis pipelines are saved in the YAML format, and the results, in the NWB format. Users with modest experience in coding can add their own analytical tools as modules written in Python. The installation procedure is described in Supplemental File 1.

### Architecture

The configuration of OptiNiSt allows researchers to easily install and test new algorithms and analytical methods. The backend uses Snakemake (https://snakemake.github.io; [16]) to control Python environments and execution flow of scripts, enabling the implementation of different analyses that require different environments to be used in the same workflow. React (https://react.dev) and FastAPI (https://fastapi.tiangolo.com) were adopted as the web library and framework for the user interface, making it a cross-platform application that functions regardless of the operating system (Fig 1).

**Fig 1.**
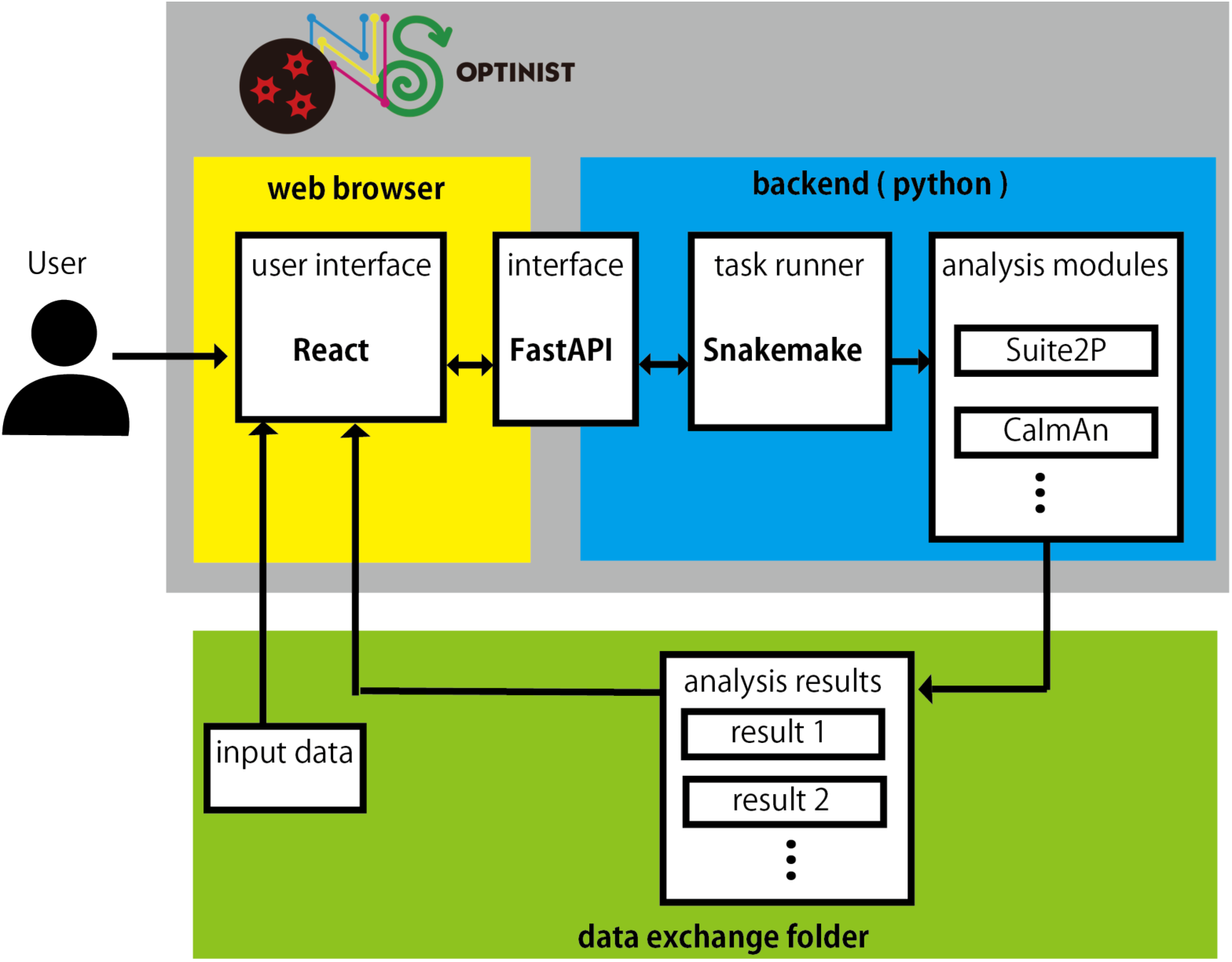
Architecture of OptiNiSt. Users interact with OptiNiSt through a web browser. According to the user’s input, it creates the SnakeMake script that controls the flow of Python scripts and runs them. The results are inspected continuously and updated, thus making them visible from the browser.

### GUI for analysis workflow design

OptiNiSt adopts a graphic interface akin to MATLAB’s Simulink or LabVIEW for users to construct analysis pipelines by connecting nodes representing functions or operations. These pipelines allow branching, thereby facilitating complex processes, and enable complete reproducibility of computational results by generating unique IDs for each operation. Such specifications enable users to create highly adaptable workflows tailored to their needs. This design also permits users to simultaneously test multiple algorithms and their parameters within the same workflow, facilitating comparison and enabling a fairer selection of algorithms.

The user creates analysis pipelines on the WORKFLOW page of the GUI (Fig 2). A single algorithm or operation module is expressed as a rectangular node. The connectors of a node (colored small rectangles) indicate the function’s input (left side) or output (right side), determining the data type by their colors (Fig 2A: 1). In the example node (caiman cnmf), the red input node indicates image data (vertical pixels x horizontal pixels x frames), and the orange output node, fluorescence data (number of cells x time). The connector type pops out when the cursor is close to it so that the user can easily confirm (Fig 2A: 2). The parameters are shown by clicking the PARAM button on the node, and the user can easily change them on the GUI (Fig 2A: 3).

**Fig 2.**
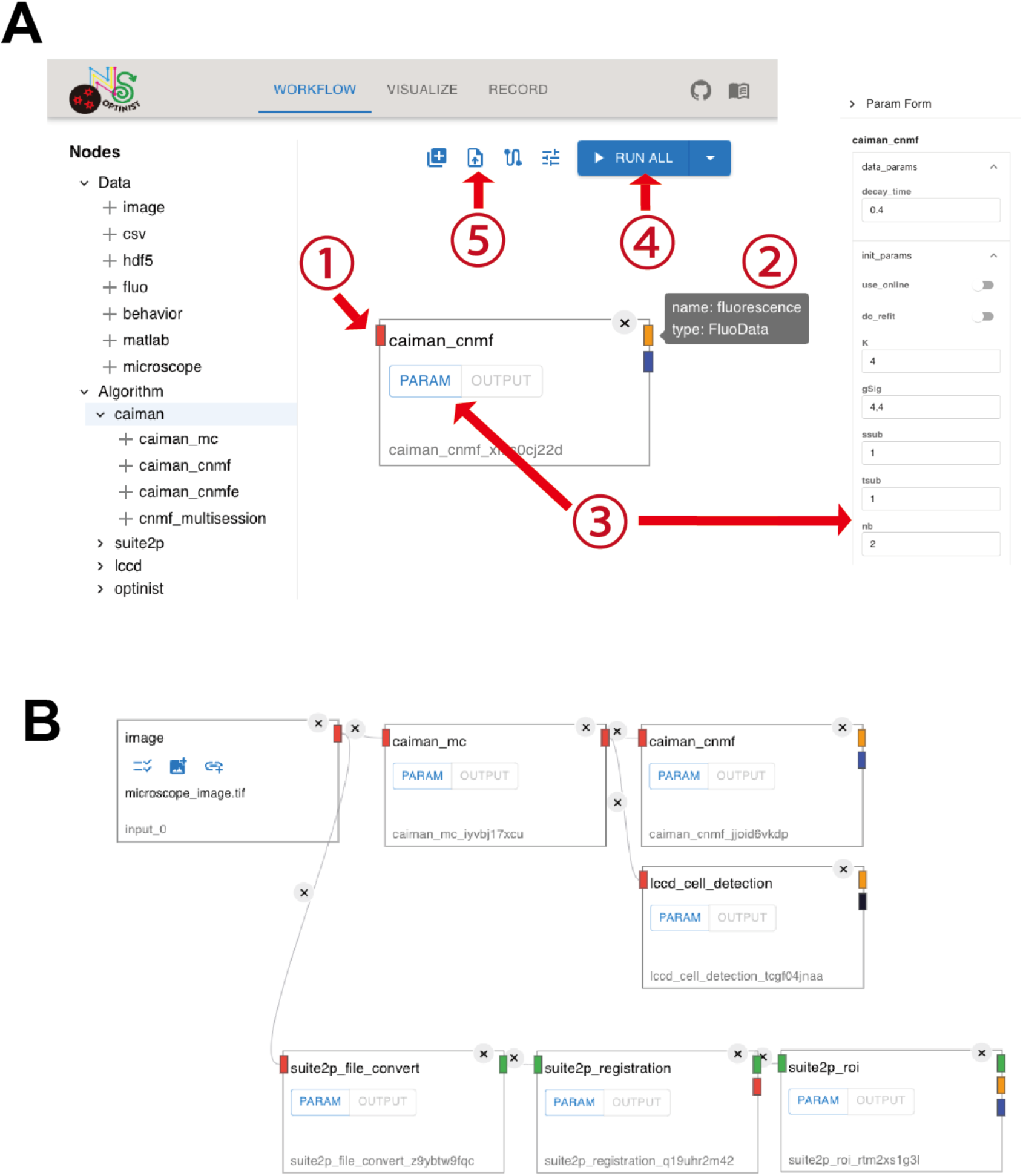
Graphical user interface for workflow. A: The appearance of the WORKFLOW page. The list on the left side shows the available operations. Clicking on the + mark introduces the node to the workspace. Each operation is represented by a rectangular node, with small connectors that assign the inputs and outputs present on the left and right sides of the node (1). Information regarding each connector can be visualized by moving the cursor over the connector (2). The parameters for an operation are shown by clicking on the PARAM button on the node (3). The RUN ALL button starts the execution of the pipeline (4). The four blue icons on the upper right are for “creating a new workflow,” “importing workflow,” (5) “editing snake make settings,” and “editing NWB settings,” respectively. B: An example pipeline that performs motion correction and cell detection using different algorithms for comparison.

The nodes are placed and connected intuitively by clicking and dragging. The rule ensuring that connectors of the same color can only be connected to each other prevents confusing the data type. The first input data can be multi-page images (.tiff) as raw imaging data, tabular data (.csv), metadata (.nwb or .h5), or MATLAB data (.mat) for other types of data, such as behavioral data or fluorescence change values of the cells. The variation in accepted datasets makes the possible pipelines flexible. A user can start with motion correction and region of interest (ROI) detection algorithms or an analysis of the population activity in already identified cells based on fluorescence change. The GUI includes a node to import data acquired from commonly used microscopes. Therefore, it is also possible to directly input raw data acquired from microscopes (Inscopix [.isxd], NIKON [.nd2], and Olympus [.oir]).

To run the workflow, the user clicks the run button on the top right (Fig 2A: 4). An indicator shows whether the process was completed with or without errors, and in case of errors, the error messages are visible when the cursor hovers over the indicator. All processing results are saved in a format that is compliant with the Optophysiology module of NWB. Metadata, such as subject IDs, data acquisition framerates, and fluorescence types, can also be input via the GUI. This feature simplifies the sometimes tedious creation of .nwb files for the purpose of generating public data. The workflow created by OptiNiSt is saved in a YAML format and can be reproduced on the working field by one click (Fig 2A: 5). Fig 2B shows an example of a workflow comparing multiple cell detection algorithms. Individual algorithms need a specific Python environment, but these environments are selected automatically while running the workflow. Upstream analyses of imaging data, such as motion correction or ROI detection, often take time. OptiNiSt has the functionality to check each step of the pipeline and execute calculations from newly added nodes or nodes with changed parameters, thereby automatically avoiding unnecessary repetition of calculations.

Currently, the cell detection algorithms implemented in OptiNiSt are Suite2P [2], CaImAn [3], and LCCD [4]. OptiNiSt also includes several pre-installed basic analytical tools (such as neural population analysis, dimensionality reduction, finding subpopulations, and neural decoding based on often-used Python modules, such as scikit-learn or statsmodels). The list of pre-installed modules is presented in Table 1. User-defined custom-made functions can be added by coding with Python.

**Table 1.**
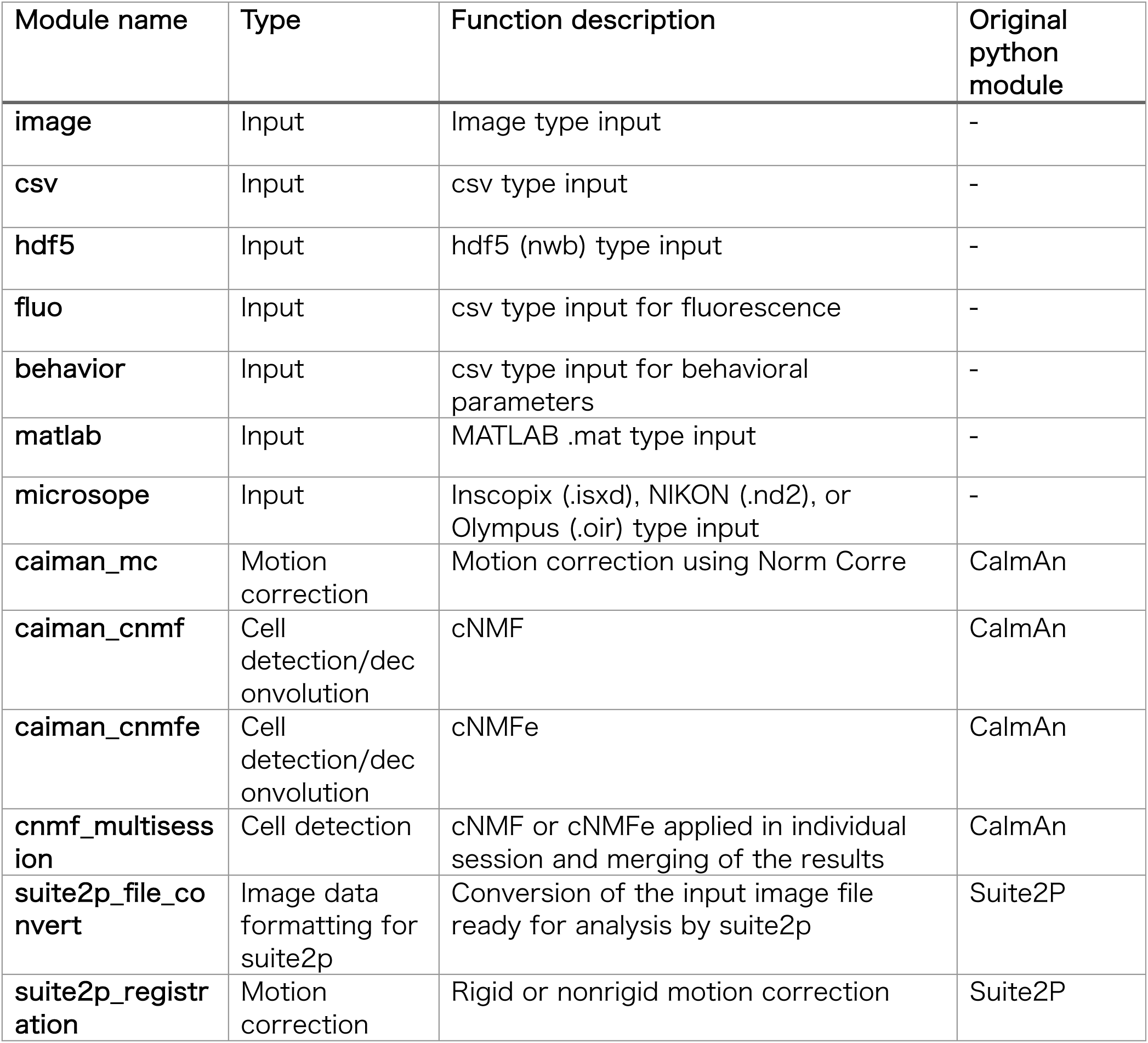
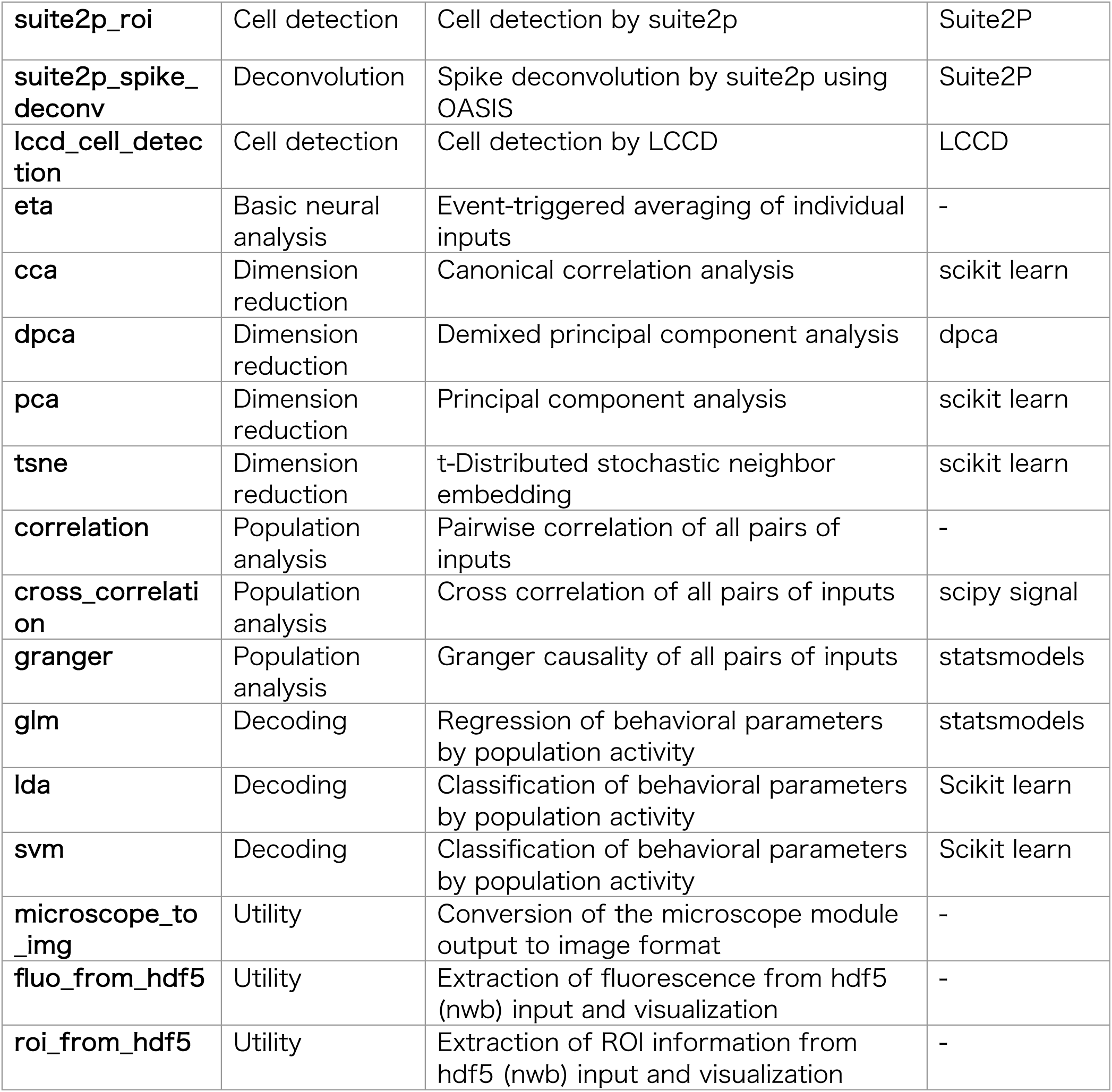
Description of pre-installed modules.

### GUI for visualization

The VISUALIZE page (Fig 3A) serves to quickly check the input images or visualize the results of analyses run on the WORKFLOW page. The VISUALIZE page allows visualization of multiple plots and comparisons. Fig 3A presents the panel for input images shown as a time-lapse video. The user selects the data and frame number to be shown in sequence. One troublesome part that needs attention in calcium imaging data analysis is checking whether the algorithm appropriately captures the cell areas. Fig 3B shows the panels for visualizing the results of cell detection with the cell area on the left and corresponding fluorescence time courses on the right. These two panels are linked to each other, and the correspondence between the cells on the map and traces is easily detected. In addition to the functionality of showing the cell area detected by algorithms, OptiNiSt allows for manually manipulating the cell area by merging, deleting, and adding new candidate cell areas and their time course (Fig 3C). The manipulation information is saved in an NWB file.

**Fig 3.**
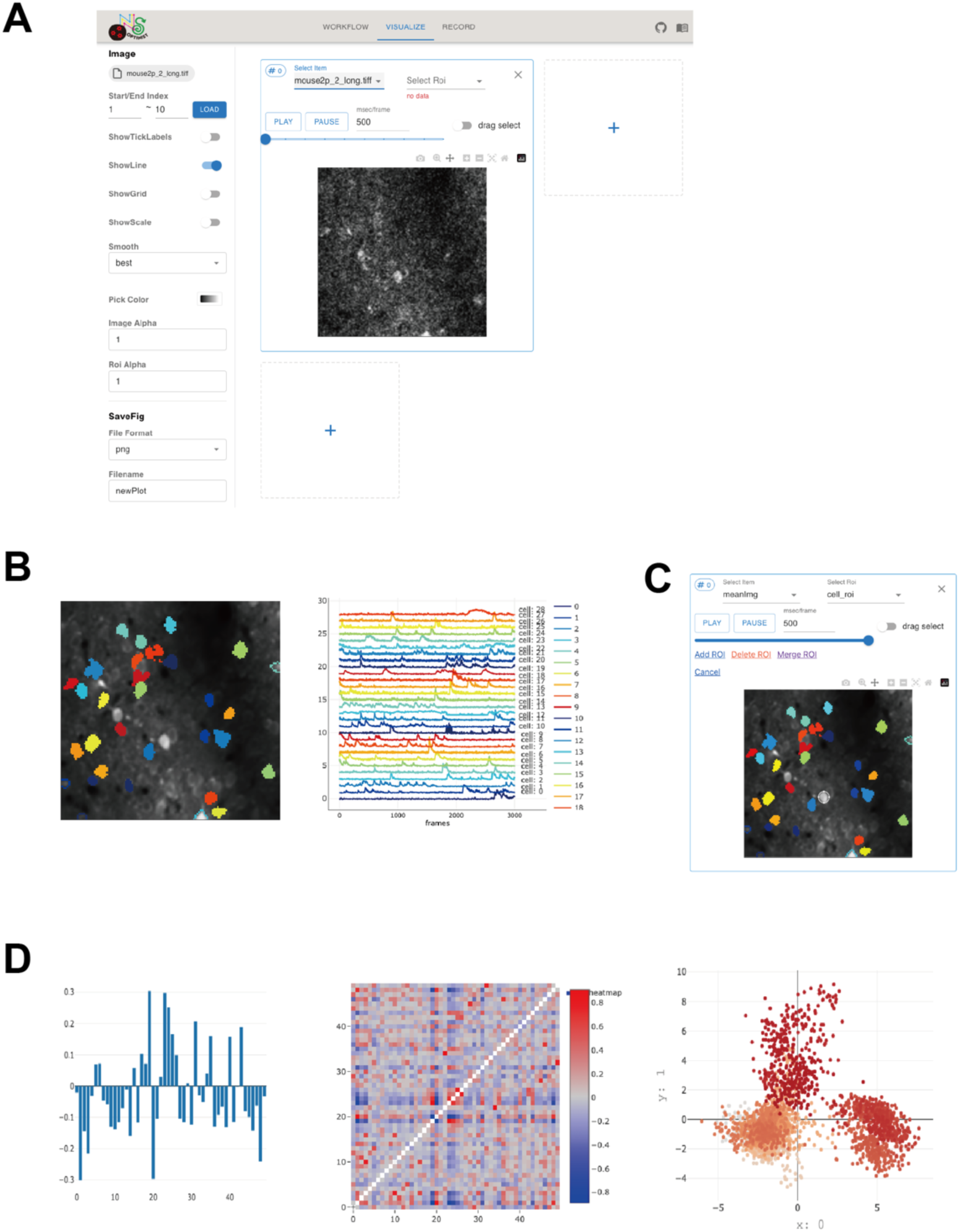
Graphical user interface for visualization. A: The appearance of the VISUALIZE page. The left side displays options available for creating plots. Here, the input image files are shown, which can be viewed as videos. By clicking the “+” sign visible on the right or bottom side, it is possible to create multiple figures. B: Figures of cell detection results. The cell area (left) and fluorescence (right) panels are synchronized. It is possible to check the cell area and corresponding fluorescence time course easily. C: Adding, merging, and deleting the cell areas and corresponding fluorescence time course are also possible. On adding cell areas, the fluorescence of the area represented by the white ellipse is added to the list of detected cells. D: Examples of the implemented plot types.

Downstream analysis applied to the detected multiple-cell activities might need other plotting styles. OptiNiSt has adopted Plotly for drawing purposes. Graphs, such as bar graphs, scattergrams, and pseudo-colored images, have already been implemented.

### GUI for workflow management

Data analysis often involves trial and error, where methods and parameters are adjusted iteratively. To facilitate the organization of created pipelines, OptiNiSt includes a mechanism for managing pipelines. The RECORD page displays a list of created pipelines. The purpose of the RECORD page is to organize the multiple pipelines, thereby minimizing confusion. This list allows the naming of each pipeline in addition to assigning unique IDs to each pipeline and facilitates differentiation by processing date, time, and duration.

The listed pipelines are easily reproduced with one click, ready to be rerun on the WORKFLOW page and to review the resulting plots. Workflow details, including each function used and indicators regarding whether it ran with or without errors, are also available. The page also has a function to save the resulting files to the local folder (Fig 4). Thus, the analysis pipeline can be easily shared with other researchers. This feature provides a very convenient option in cases in which it is necessary to publicly release all analysis programs, such as to meet the requirements of journals.

**Fig 4.**
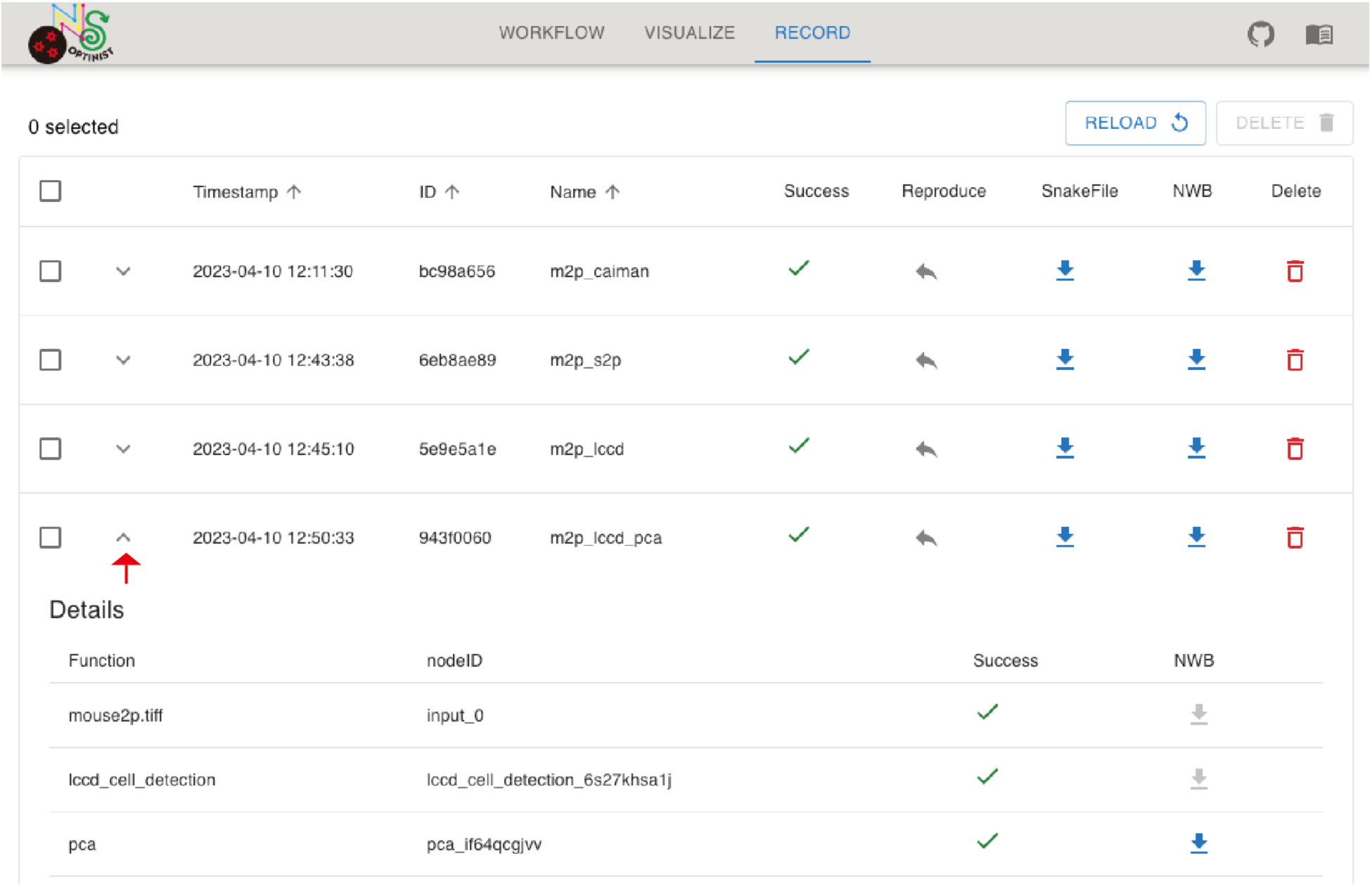
Graphical user interface for recording. When a pipeline is created, it is automatically registered in this list. The ID is unique to each pipeline. The name is assigned by the user. When the pipeline is completed without errors, a check is placed in the Success column. By clicking the Reproduce button, the pipeline is recreated on the WORKFLOW page. Snakefiles and NWB files can also be downloaded with one click. A pipeline is made up of multiple functions. Clicking on the part with the red arrow displays the details.

### Saving format

OptiNiSt describes the details of the workflow in YAML format, which is crucial for pipeline reproducibility. From this file, it is possible to generate a Snakemake script, which is required for actual computations (also in YAML format). The computation results are documented in HDF format files compliant with NWB guidelines. In addition to commonly handled variables in calcium imaging, such as fluorescence time course, cell position, and cell ID, this file simultaneously saves outputs in the same file if the original function supports saving NWB files. For motion correction results, in which the image file size tends to be large, users have the option to choose whether or not to save, providing flexibility tailored to their demands. Therefore, with these files, the workflow and computation results are fully documented.

### Implementation of custom-made functions

The OptiNiSt is designed such that users can add their own analysis functions as modules. Fig 5 shows an example of inserting a new algorithm and its function. Users only need to add a limited number of files to the appropriate directories and modify one of the existing files. The files to be added include 1) the main function file (Fig 5A) that includes the main contents of the function, 2) initialization file that initializes the module and determines the hierarchy of the modules (Fig 5B), and 3) the YAML file that defines the parameters and their default values related to the new function (Fig 5C) and conda environments (Fig 5D). Additionally, to make the function usable on the GUI, the information on the new function module should be added to the existing initialization file (Fig 5E).

**Fig 5.**
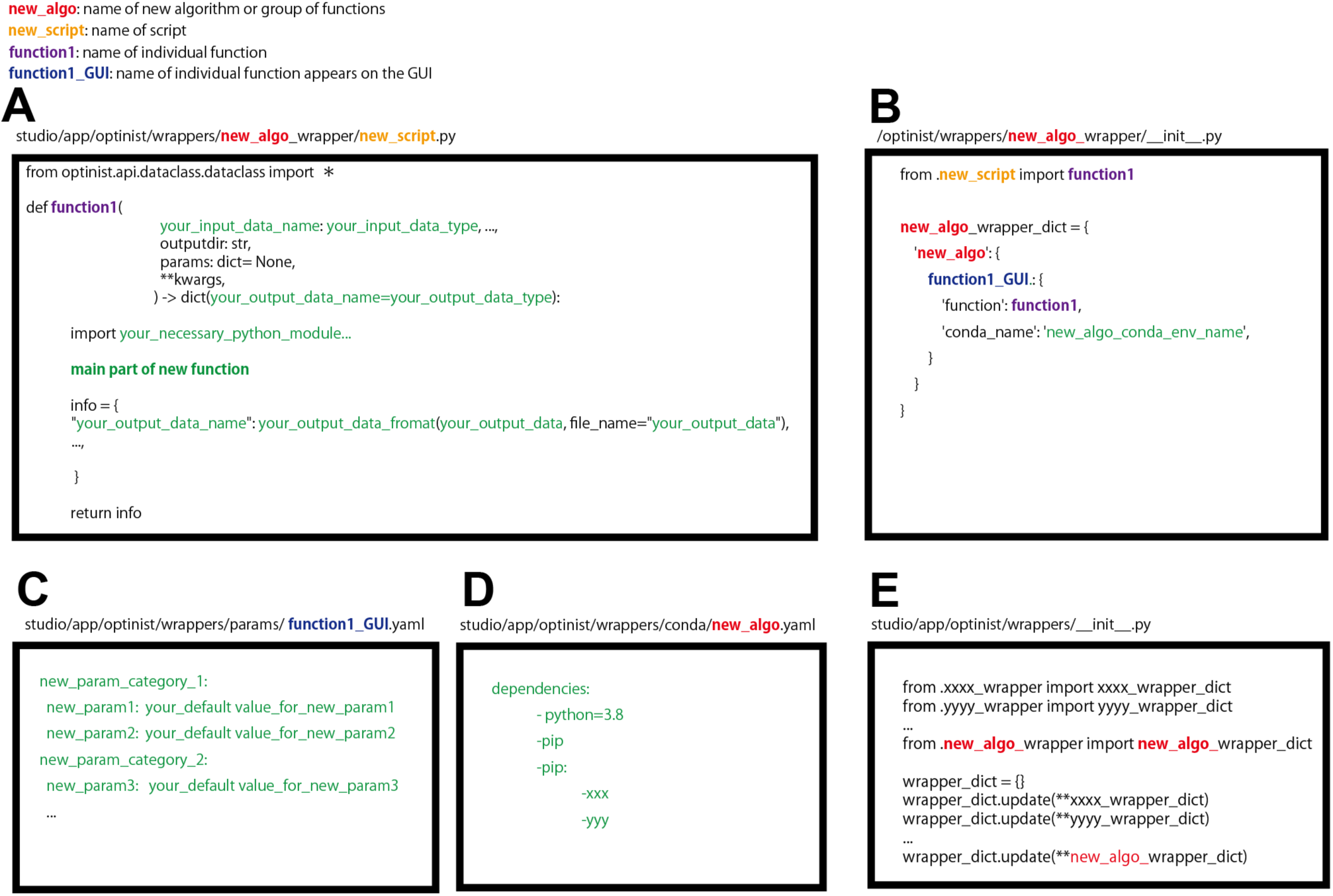
Adding a new function module. Files should be placed in the appropriate place, which is shown on the top of each panel. A: The main function determining the calculation should be written in a fixed format, enabling easy input and output configuration. Green letters indicate the lines that the user defines. B: The initialization file for the module. C-D: The parameters and conda environments are defined in YAML style. E: To make the module appear in the graphical user interface, information on the new function module should be added to the existing initialization file.

### Script execution

OptiNiSt provides an option to execute the workflow designed by GUI on computing servers by downloading the Snakemake config file, which includes all the information about the workflow, from the RECORD page. Input and output paths are changeable by environmental variables for OptiNiSt. The script, run_cluster.py, executes the workflow with an option assigning the Snakemake config file path. This procedure simply runs backend processes and is convenient for executing pipelines for multiple large datasets on HPC or clusters.

## Results

Here we present examples of building workflows and visualizing the analysis results using sample 2-photon imaging data obtained from the parietal area of a mouse during auditory stimulation from 12 speakers surrounding the animal [17]. The details of the data are described in Supplemental File 1.

### Comparison of motion correction algorithms

The first step of imaging data processing is motion correction. The analysis pipeline shown in Fig 6 compares the performance of the following motion correction algorithms: caiman motion correction (Norm Corre: [18], TurboReg [19], and Suite2P rigid and nonrigid registrations. Fig 6A shows the workflow. To compare the same image data, branching from the image node allows input to each motion correction algorithm. OptiNiSt seamlessly handles such branching processes. While comparisons of different algorithms are necessary, the environment is automatically segmented for each node. Clicking on OUTPUT in each node displays the mean image after performing motion correction (Fig 6B), enabling quick comparison.

**Fig 6.**
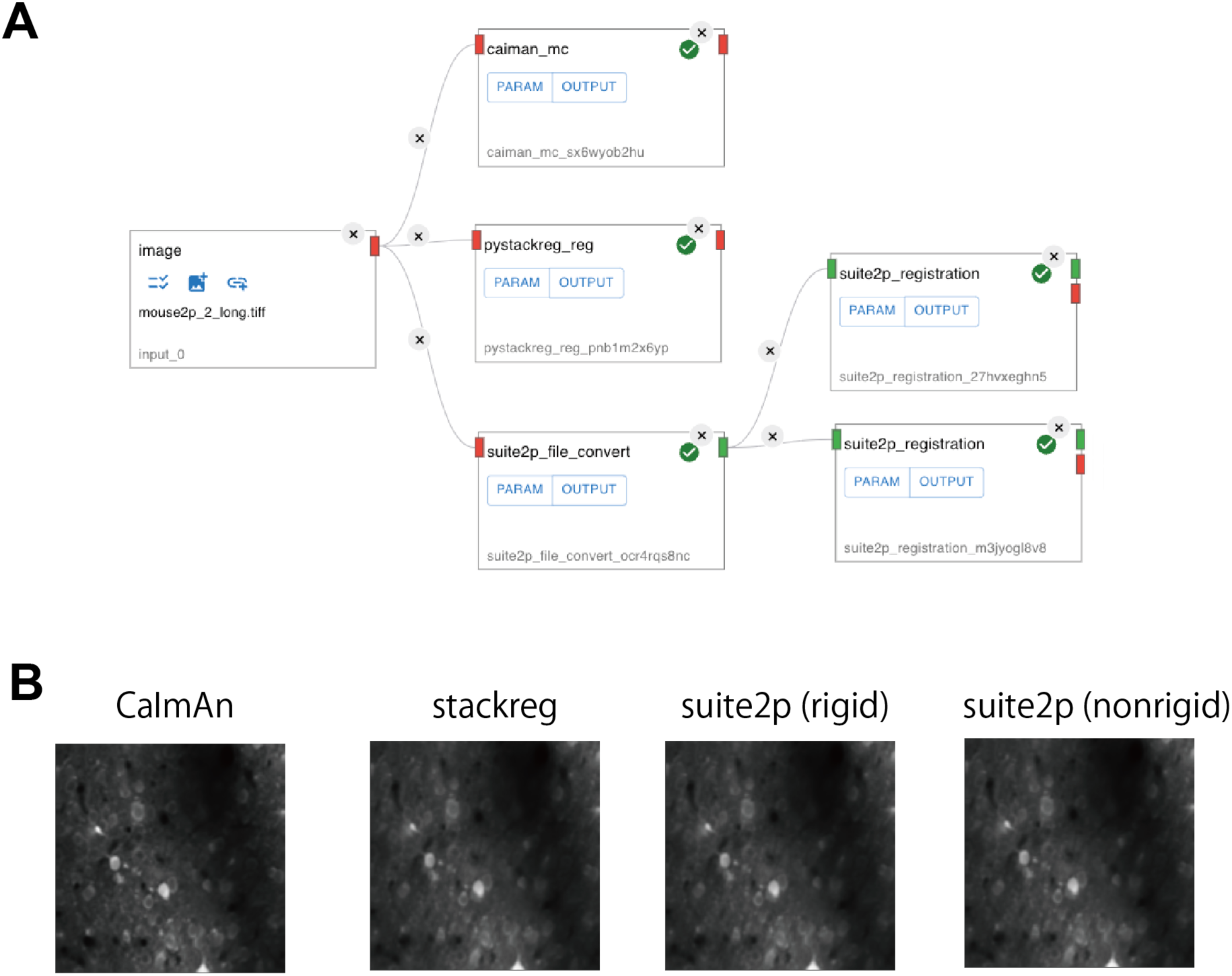
Example of motion correction comparison. A: The pipeline to compare different motion correction algorithms, B: The mean (across frames) images for individual algorithms

### Comparison of cell detection algorithms

The next demonstration (Fig 7) compares different cell detection algorithms. The data used here are the same 2-photon data used in Fig 6. In this example, we compared cell extraction algorithms: CaImAn cnmf [3], Suite2P [2], and LCCD [4]. The number of cells extracted by each algorithm was 29, 55, and 28, respectively. These numbers can change depending on parameters. Here, the default parameters were used. Fig 7A shows the pipeline for comparison. The raw TIFF file was motion corrected using NormCorre provided by CaImAn, and the output of motion correction was bifurcated and input to each cell detection algorithm. The resulting cell areas (Fig 7B) and corresponding fluorescence time course (Fig 7C) are shown on the VISUALIZE page. By linking the cell ROI and fluorescence panels, the correspondence between cell area and their fluorescence can be checked easily. The fluorescence time course of cells indicated by the white arrows in Fig 7B is shown as the upper three traces in Fig 7C (red rectangles). These tree cells were detected across all three algorithms, but the resulting traces were not the same; specifically, the top cells of cNMF looked quite different from those of the other two.

**Fig 7.**
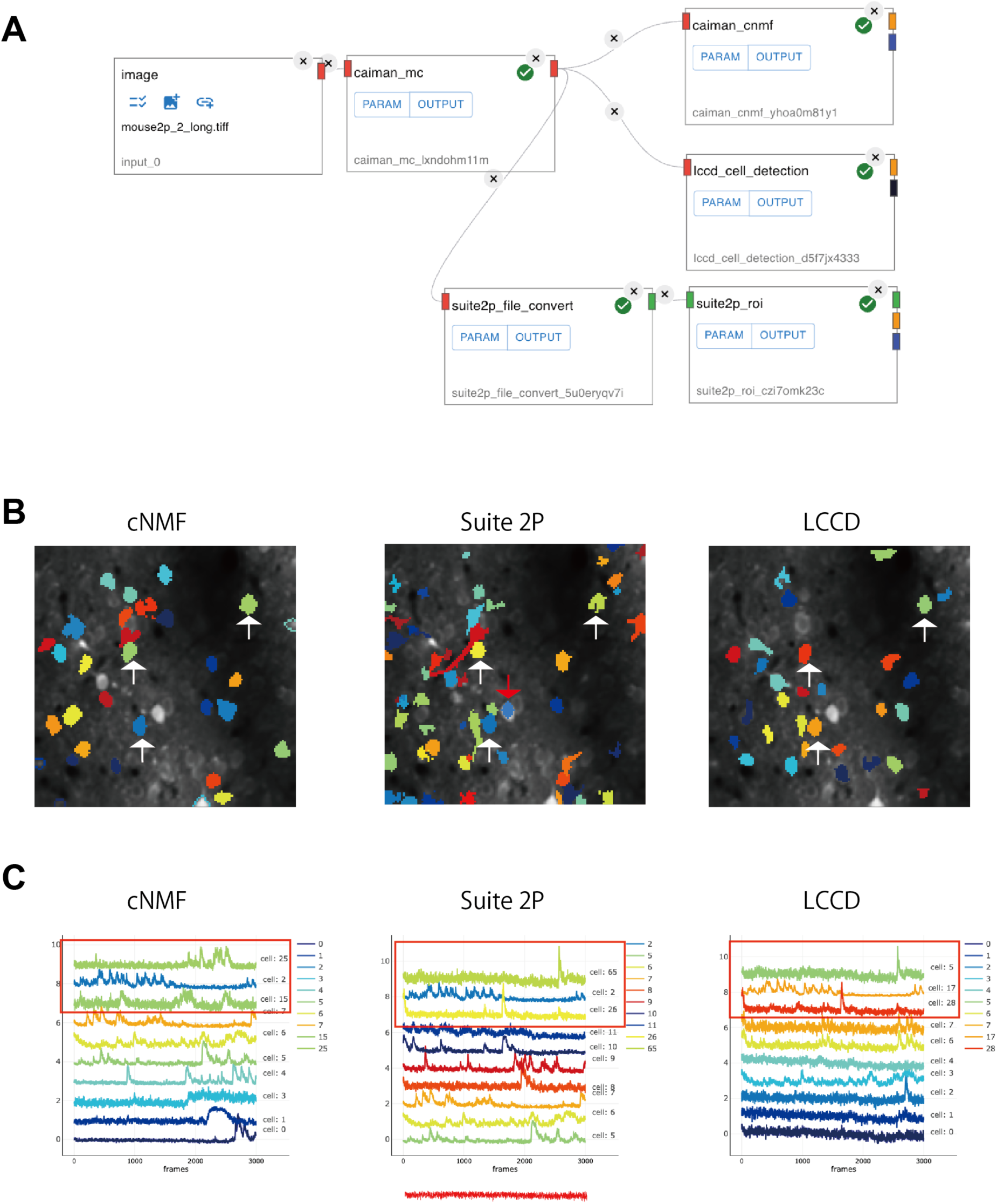
Example of cell detection comparison. A: The pipeline to compare different cell detection algorithms. B: Cell areas detected by three different algorithms. Three example cells detected by all the algorithms, as indicated by white arrows. C: The fluorescence time course of detected cells. The top three traces indicated by red rectangles correspond to the cells indicated by the arrows in B.

### Neural population data analyses

OptiNiSt supports downstream analyses of neural population activity data, as exemplified in Fig 8. Here, the output of suite2p (fluorescence time course, iscell, and cell position), which was saved in the previous example (Fig 7) as a .nwb file, was fed into three different analytical tools: pairwise correlation, event-triggered averaging, and CEBRA [20] (Fig 8A). The number of ROIs detected by suite2p was 266, and 55 cells were qualified among them. Information on whether an ROI has passed the qualification (iscell), the fluorescence of individual cell, and data on cell areas are saved in single .nwb file and the three different input modules were used to assign each of them.

**Fig 8.**
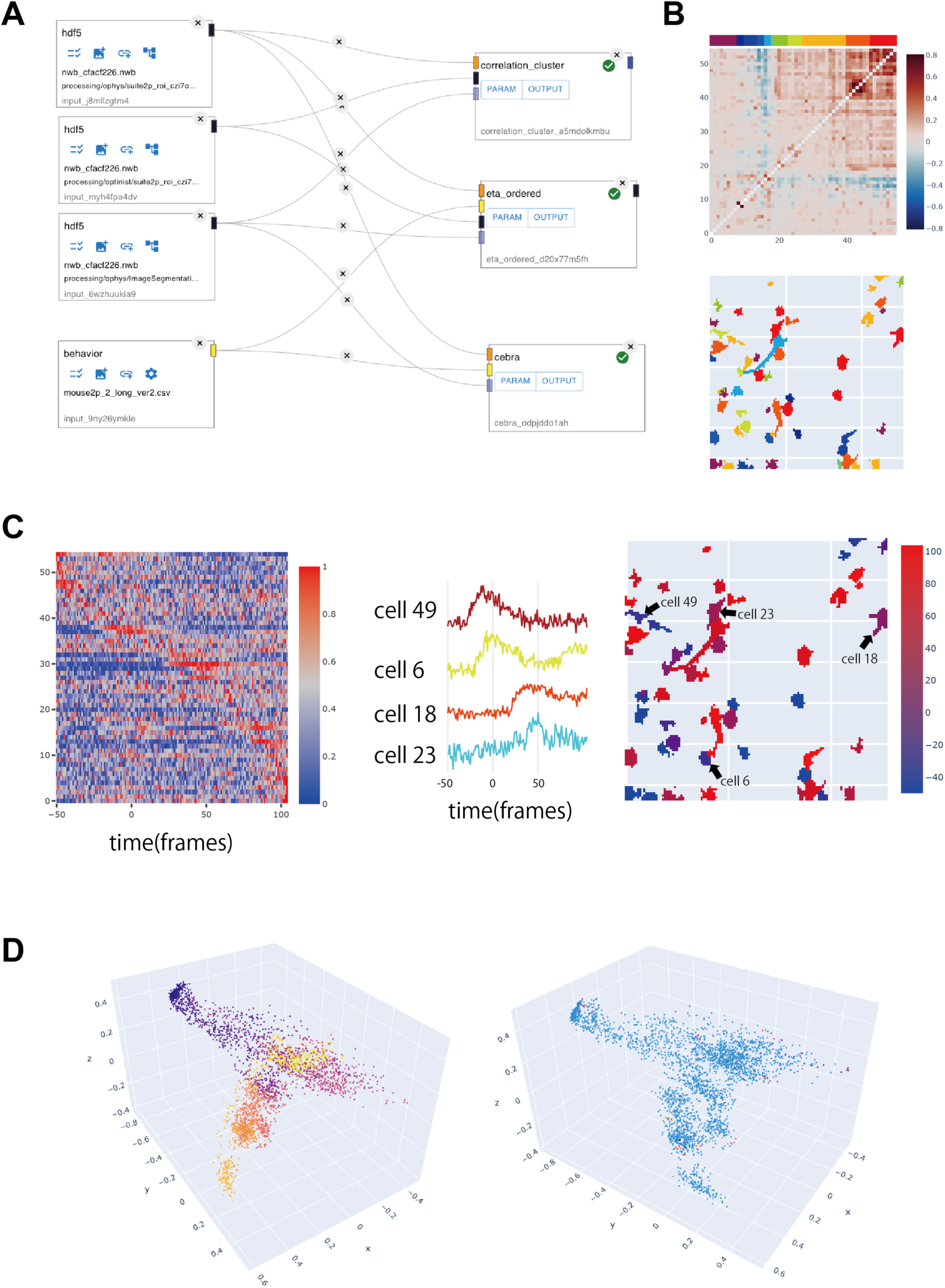
Example of downstream analysis. A: The pipeline for performing pairwise correlation analysis, triggered averaging, and CEBRA. B: Correlation matrix ordered by hierarchical clustering (upper). The colored bar on the top indicates the cluster. Cell areas are indicated and colored based on the cluster (lower). C: Stimulus-triggered averaging of mean fluorescence across trials (left). The example mean traces showing earlier and later increases in activity (center). Cell areas are indicated and colored based on the peak time (right). The cells indicated by the arrows are the same as those in the center trace. D: Latent values identified using multiple neuronal activity, stimulus, and licking information using CEBRA. The colors of the images are based on licking (left) and stimulus (right), respectively.

The module, “correlation_cluster,” calculates the pairwise Pearson’s correlation coefficients between all neurons and clusters based on hierarchical clustering. The threshold for clustering is set to 0.5 here. The resulting correlation matrix ordered by clusters is shown in the upper panel of Fig 8B. In this example, two clusters indicated by the color red and orange showed strong correlations among cells. The positions of these functionally clustered cells can be examined as shown in the lower panel of Fig 8B, which depicts cluster identity by color.

The node “eta_ordered” calculates the event-triggered averaging of the fluorescence of an individual cell. The trigger time is based on the behavior data, which are imported from a .csv file. Data length should be the same as the frame number. In this example, the trigger is the timing of auditory stimulus onset. Fig 8C (left) shows the resulting activity. Some cells show a peak around frames 0 and 50. Examples of these cells are shown in the middle panel. The position of these cells is shown in the right panels, together with the peak time indicated by different colors.

The combination of relatively straightforward analyses like these is essential for checking the quality of recorded data and observing overall response trends, but coding them from scratch can be cumbersome. In OptiNiSt, it is possible to embed specific calculations for calcium imaging into nodes. These nodes allow customization in terms of saving values to NWB and drawing figures with relatively minimal coding.

To demonstrate the possibility of creating nodes for relatively advanced calculations, a node for “CEBRA” [20], which finds latent space based on multiple neural activity and behavioral parameters, was introduced. As this example is for demonstration, we only show the latent space discovered by supervised learning. In this example, the licking behavior and stimulus timing from 12 different positions, as well as the fluorescence of multiple cells, were fed into the node to find the 10-dimensional latent activity. The figure shows three dimensions chosen arbitrarily from 10 with the behavioral parameters as colored codes.

### Performance in large-scale datasets

Motion correction and cell detection may require significant memory size and computation time, depending on the specific algorithms used and availability of GPUs. The OptiNiSt framework itself does not have any constraints for the execution of large datasets. As an example, it has been confirmed that the modules pre-installed in OptiNiSt can analyze 12 GB of imaging data in one go using a Desktop PC (Mac Pro, OS: Ventura, memory size: 192 GB). Additionally, OptiNiSt incorporates both the multi-session cNMF, which performs cell detection on multiple small datasets, and the online cNMF, which iteratively analyzes data while loading them.

## Availability and Future Directions

OptiNiSt is an open-source project, which is anonymously available on a source repository hosted by GitHub (https://github.com/oist/optinist). Users can install OptiNiSt to Linux, Mac, and Windows machines by the Python “pip install” command or by downloading a virtual machine from DockerHub (oistncu/optinist). The user guide, documentation, and tutorials are provided at https://optinist.readthedocs.io/en/latest/. There are Slack user communities available to discuss the use of OptiNiSt (https://oist.enterprise.slack.com/archives/C02V1FP6Z8W), and there is a possibility of discussion on GitHub.

The future direction includes the extension of the available analysis modules by making it easy for researchers to implement their methods as OptiNiSt modules. We hope that the OptiNiSt framework can promote testing of various analytical methods, contributing to the creation of reproducible analysis pipelines.

## Acknowledgments

We are grateful to the developers of the cell detection algorithms—suite2p, CaImAn, and LCCD—for making them open source. We thank the software engineers, Shogo Akiyama, Rei Hashimoto and Yuta Fukuda for their exceptional technical support and development of the software architecture. We are grateful for the help and support provided by the Scientific Computing and Data Analysis section of the Research Support Division at OIST and for the English language editing by Editage (www.editage.jp). We thank Yukiko Yamane for creating the logo.

## Notes

https://optinist.readthedocs.io/en/latest/

https://github.com/oist/optinist

https://doi.org/10.5281/zenodo.13357961

### Competing Interest Statement

The authors have declared no competing interest.

