## Supplemental text for "Optical Neuroimage Studio (OptiNiSt): intuitive, scalable, extendable framework for optical neuroimage data analysis"

### Supporting information

#### Installation (general procedure)

Docker image and pip installation methods are provided in addition to Open codes on GitHub. Installation for Windows, Mac, and Linux is supported. We have provided two different setting modes: standalone and multiuser. The standalone mode assumes that OptiNiSt is installed on a PC and used by a single user. The multiuser mode allows authentication and use by multiple users and running of multiple workflows simultaneously, providing a single platform for using a high performance server machine for individual or collaborative analyses. OptiNiSt uses mysql or mariadb for the management of accounts, and Firebase Authentication, for authentication. Installation of both modes is simple, and the procedure is documented (https://optinist.readthedocs.io/en/latest/).

#### Dataset for the examples

We used 2-photon imaging data obtained from the parietal area of a mouse during auditory stimulation from 12 speakers surrounding the animal. We prepared cropped small size imaging data (128 pixels x 128 pixels x 3000 frames, 100 MB) and behavior data for the demonstration, and the large size original data (512 x 512 x 22500, 12 GB) were used for confirming that OptiNiSt can handle an original-size dataset. Both datasets are available at Zenodo repository.
Zenodo repository: https://doi.org/10.5281/zenodo.13357961.

#### I Instruction for reproducing the analyses

To reproduce the analysis pipeline shown in Fig. 7, any installation method suitable for general users (e.g., Docker or pip install) or for developers (e.g., downloading the repository from

GitHub) can be used. The installation instructions are documented in https://optinist.readthedocs.io/en/latest/installation/index.html. To reproduce the analysis pipelines shown in Figs. 6 and 8, an additional user-defined module needs to be added. For this purpose, installation by downloading the repository from GitHub is preferrable. By adding the "wrappers/paper" folder from the Zenodo repository to the "optinist/studio/app/optimist/wrappers" folder in the cloned OptiNiSt repository and replacing __init__.py in the same folder with the version from the Zenodo repository, the additional module will become available. In addition, the software with additional user-defined module (optinist-main) is archived on the Zenodo repository.

Reproducing Fig. 6: Figure6_workflow.yaml can be uploaded by clicking on the import icon in the WORKFLOW field (see Fig. 2A: 5). Before running the pipeline, the data (mouse2p_2_long.tiff) should be assigned by clicking on the input module and selecting it.

Reproducing Fig. 7: Figure7_workflow.yaml and the data (mouse2p_2_long.tiff) need to be uploaded and selected as Fig. 6.

Reproducing Fig. 8: Figure8_workflow.yaml needs to be uploaded as Fig. 6. The three input hdf5 input modules should assign the same file, nwb_cfacf226.nwb. These three modules assign different content in the same file. For the top hdf5 module which is connected to the fluorescence (orange) connectors, processing/ophys/suite2p_roi_czi7omk23c /Fluorescence/data in nwb_cfacf226.nwb should be assigned. For the middle hdf5 module which is connected to the black connectors, processing/ophys/optinist/suite2p_roi_czi7omk23c_all_roi_img/data in nwb_cfacf226.nwb should be assigned. For the bottom module which is connected to the iscell (blue) connectors, processing/ophys/ImageSegmentation/suite2p_roi_czi7omk23c/iscell should be assigned. The behavior input module should assign mouse2p_2_long_with_headder.csv
